# Decoupling of Dopamine Release and Neural Activity in Major Depressive Disorder during Reward Processing Assessed by Simultaneous fPET-fMRI

**DOI:** 10.1101/861534

**Authors:** Xue Zhang, Fuyixue Wang, J. Paul Hamilton, Matthew D. Sacchet, Jingyuan Chen, Mehdi Khalighi, Ian H. Gotlib, Gary H. Glover

**Affiliations:** Center for Biomedical Imaging Research, Department of Biomedical Engineering, School of Medicine, Tsinghua University, Beijing, China; Radiological Sciences Laboratory, Department of Radiology, Stanford University, Stanford CA, United States; Harvard-MIT Health Sciences and Technology, MIT, Cambridge, MA, United States; A. A. Martinos Center for Biomedical Imaging, Department of Radiology, Massachusetts General Hospital, Charlestown, MA, United States; Center for Social and Affective Neuroscience, Linköping University, Sweden; Center for Depression, Anxiety, and Stress Research at McLean Hospital, Harvard Medical School, Boston, MA, USA; Department of Radiology, Harvard Medical School, Boston, MA, USA; Department of Psychology, Stanford University, Stanford CA, United States

**Keywords:** fPET-fMRI, dynamic PET analysis, GLM, MDD, reward processing

## Abstract

The interaction of the midbrain dopaminergic system and the striatum is implicated in reward processing; it is still unknown, however, how this interaction is altered in Major Depressive Disorder (MDD). In the current study, we related the dopamine release/binding inferred by [^11^C] Raclopride functional Positron Emission Tomography (fPET) to neural activity monitored by blood-oxygen-level-dependent (BOLD) functional magnetic resonance imaging (fMRI) in adults diagnosed with MDD and healthy controls (CTL). Participants completed a monetary incentive delay (MID) task during simultaneous [^11^C] Raclopride fPET and fMRI. Instead of the usual kinetic modeling method for analyzing dynamic PET time activity curves (TACs), we used a simpler general linear model (GLM) approach, which includes introducing a fPET dopamine activation response function to model changes in the TAC associated with the MID task. In addition, using simulations, we show that the GLM approach has several advantages over kinetic modeling. This is achieved without invoking erroneous steady-state assumptions or selecting a suitable reference region. Our results include the observation of both decreased fMRI activation and dopamine release/binding in the striatum in the MDD cohort, implying a reduced reward processing capacity in MDD. Furthermore, in the MDD group, individuals with lower fMRI activations in the right middle putamen and ventral medial prefrontal cortex (vmPFC) had higher reflection rumination scores, and individuals with lower dopamine release/binding in the left putamen and the right nucleus accumbens (NAcc) also had higher reflection rumination scores. Significant cross-modal inter-subject and intra-subject correlations of dopamine release/binding and fMRI activation were observed in the CTL group, but not in the MDD group. The intra-subject correlation of the two modalities was negatively associated with reflection rumination scores in the CTL group, indicating that decoupling of the dopaminergic system and striatum may be important in the pathophysiology of MDD.

## 1. Introduction

Functional magnetic resonance imaging (fMRI) has relatively high temporal and spatial resolution but can have modest sensitivity and specificity because it measures the blood oxygen level change caused by neural firing instead of the neural activity itself (Ekstrom, 2010; Glover, 2011; Sejnowski et al., 2014). In contrast, positron emission tomography (PET) has high imaging sensitivity and specificity because it images targeted exogenous radiotracers. Raclopride is an antagonist of the dopamine D_2_ receptors which are primarily expressed in the striatum (Elsinga et al., 2006). [^11^C] Raclopride PET can be used to image dopamine release and binding at D_2_ receptors because dopamine release displaces Raclopride, thereby causing a PET signal decrease. However, the temporal and spatial resolution of PET are limited relative to fMRI due to low signal to noise ratio (SNR) and lengthy tracer binding dynamics. Recent technical advances in PET (such as the time of flight technique) enable dynamic functional PET (fPET) image acquisition with 15 s to 1 min temporal resolution while maintaining an acceptable SNR (Surti, 2015). With the use of simultaneous [^11^C] Raclopride fPET-fMRI (Elsinga et al., 2006; Judenhofer et al., 2008), researchers can dynamically track the coupling or interaction of whole-brain neural activity and dopamine release/binding at D_2_ receptors, thus yielding deeper insights about brain function under normal and disease conditions.

The ventral tegmental area (VTA), a primary node of the dopaminergic system, sends dopaminergic neural projections to numerous brain regions ranging from cortex to subcortical structures (Beier et al., 2015). Functional interactions between the midbrain dopaminergic system and the striatum, and especially the nucleus accumbens (NAcc), are a crucial component of reward processing (Haber and Knutson, 2010). Dysfunction of the dopaminergic system has been linked with the pathophysiology of Major Depressive Disorder (MDD). Several researchers have reported reductions in fMRI activation in the striatum in MDD patients during reward anticipation (Pizzagalli et al., 2009; Takamura et al., 2017; Ubl et al., 2015b; Whitton et al., 2015), while other investigators have reported no MDD-associated differences in the striatum (Knutson et al., 2008). PET studies have implicated aberrant baseline dopamine binding potential in the striatum in MDD (Hamilton et al., 2018; Hirvonen et al., 2008a; Meyer et al., 2006; Montgomery et al., 2007a; Sun and Sun, 2017; Yatham et al., 2002), and their findings are similarly inconsistent, with reports of both increased and decreased dopamine binding. Simultaneous [^11^C] Raclopride fPET-fMRI assesses both neural activity in the reward network and dopamine release from the substantia nigra and ventral tegmentum to striatal targets; this bimodal technique may provide insight concerning reward processing in MDD and help to explain the discrepant findings of previous studies. Researchers have demonstrated using separate fPET and fMRI scans that the dopamine release/binding at D_2_ receptors is correlated with local hemodynamic changes in humans not selected for psychiatric status (Schott et al., 2008). A study using concurrent Raclopride fPET-fMRI has also confirmed coupling between the replacement of radiolabeled Raclopride and nonradiolabeled Raclopride and the BOLD fluctuations (Sander et al., 2013). This was achieved by varying non-radiolabeled exogenous ligands to imitate the activity of the endogenous neurotransmitter. However, it is still unknown whether coupling between the dopaminergic system and neural activation in the reward network persists in MDD; that is, whether altered fMRI activation in the reward network is related to deficiencies of the dopaminergic system. Recently, trait rumination has been reported to influence the anticipation phase (when dopamine releases dramatically) of reward processing among never-depressed people (Kocsel et al., 2017), however such studies are at an early stage. In the current study, we are able to explore the relation of trait rumination with the dopamine release and the fMRI activation alone, and the coupling of the two modalities to examine the neural substrates of the association of rumination and reward anticipation.

To date, the estimation of acute dopamine release/binding using dynamic fPET imaging is still being developed. The “classic” kinetic approach relies on a steady state assumption that the bound and free tracer are in an instantaneous equilibrium. An example kinetic approach includes the simplified reference region model (SRRM) that has few estimated parameters and does not require arterial blood sampling (Lammertsma and Hume, 1996a). However, inclusion of a reward processing task during PET scanning may alter tracer binding, and therefore explicitly violate the steady-state assumption. Several *dynamic* kinetic models have been proposed which include time-dependent varying parameters for the estimation of the release of dopamine under a reward task completed in a single scan (Pappata et al., 2002). These have included modeling the activation state using an exponential decay function (Alpert et al., 2003) or a gamma variate function (GVF) (Normandin et al., 2012; Sander et al., 2013). The performance of commonly used dynamic kinetic models largely depends on the selection of a reference region that is devoid of specific binding, as well as the activation function. To date, activation functions have not been widely used or validated. Given that the major challenge in revealing acute dopamine release is to eliminate potential contamination from the baseline uptake and wash-out curve, we propose here to regress out the background waveform from the time activity curve (TAC) using the general linear model (GLM). The GLM has been widely used in fMRI studies to identify the neural activation patterns induced by a specific task, wherein the modeled task responses, yielded by convolving the task paradigms with a characteristic hemodynamic response function, are included as design covariates (Friston et al., 1994). The feasibility of GLM in fPET studies has been demonstrated by 2-[^18^F]-fluorodeoxyglucose (FDG)-based studies to reveal metabolic changes during a visual task (Rischka et al., 2018; Villien et al., 2014). In the present study, we propose to extend the same framework to estimate dopamine release during a reward processing task using Raclopride fPET, the baseline uptake and wash-out curve of which have not yet been studied before. Here, we introduce a dopamine activation response function that describes the dopamine release in a GLM as an effective method to characterize tracer kinetics.

In the current study, we first used simulations to demonstrate the feasibility of using a GLM as a simpler alternative to traditional kinetic modeling for tracking the dynamic changes of dopamine release/binding. We then conducted a simultaneous [^11^C] Raclopride fPET-fMRI experiment to characterize and compare the fMRI activation and dopamine release/binding between the MDD group and healthy controls (CTL) during a monetary incentive delay (MID) task (Knutson et al., 2008). Finally, we examined relations between dopamine release and fMRI activation to probe the potential interaction between the dopaminergic system and the reward network. We hypothesized that, relative to multi-compartment kinetic models, the GLM could show comparable or improved performance in the estimation of instant dopamine release. We hypothesized further that the MDD group would demonstrate reduced activation in task fMRI, dopamine release and a disrupted coupling between the information captured by both imaging modalities, and the reduced fMRI activation, fPET activation, and the coupling between the two modality were associated with the trait rumination in the MDD.

## 2. fPET Analysis Methods

### 2.1. Theory

#### 2.1.1. Kinetic Modeling

The classic SRRM (Gunn et al., 1997; Lammertsma and Hume, 1996b) is defined by:

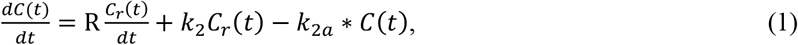

where 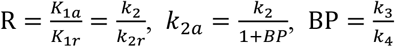, the subscripts *a* and *r* refer to the target and reference region respectively. *C*(*t*) and *C_r_*(*t*) denote the radioactivity concentration in the target region and reference region. *K*_1_ and *k*_2_ are the rate constants for the transfer between plasma and reference/free compartment, *k*_3_ and *k*_4_ are the rate constants for the transfer between free and bound compartment, while *k*_2*a*_ is the overall rate constant for transfer from the specific compartment to plasma in the target region. R is the delivery ratio of the plasma to the target region and the reference region, and BP is the binding potential of the radiotracer in the target region.

The linear extension of the simplified reference region model (LSSRM) was proposed to simulate the activation state by making the parameters time-dependent (Alpert et al., 2003) in the SRRM (Eq. 2):

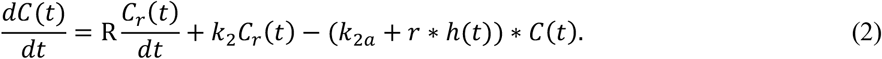

Here *r* determines the activation level, *h*(*t*) describes the change in the activation state, taken here as an exponential decay function. An operational equation was derived for convenient numeric computation by replacing integration by accumulation after integrating Eq. 2.

#### 2.1.2. GLM

A TAC example of baseline kinetics (without performing a reward task) from a preliminary study is shown in Fig. S1A. Choosing a GVF to fit the background kinetics, we have

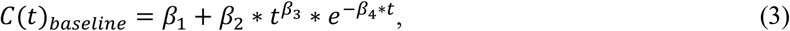

with *β*_1_ = 0, *β*_2_ = 1, *β*_3_ = 0.5, *β*_4_ = 1/40, and t represents fPET time frame number. Figure S1B shows the TAC of the same subject performing a 10-min MID task (described in section 3.3). After regressing out the baseline kinetics modeled by the GVF, the residuals exhibited a prominent modulation by the reward task – the signal increase was in good correspondence with the task timing. The residuals were fitted by an exponentially decaying impulse response function convolved with the task design (Eq. 4, Fig. S1B):

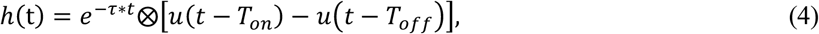

where *T_on_* is the task onset time and *T_off_* is the task end time.

We used *h*(*t*) to model the task-induced fPET signals in our experiment with *τ* = 1/3 estimated from the preliminary data (Fig. S1B). For the GLM analysis, a GVF together with the activation function *h*(*t*) were entered in the final fitting model,

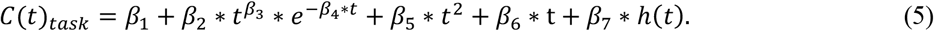

The first principal component of all TACs within the striatum was computed for each subject and taken as the baseline uptake and washout curve (background curve) of all voxels. The *β*_2_,*β*_3_,*β*_4_ parameters determining the shape of the GVF were fitted once for the first principal component and fixed for all voxels; hence the analysis is only focused on those regions with D_2_ receptors instead of the whole brain. To allow variation in the baseline uptake and washout curves of different voxels, a quadratic function was also included to compensate for potential differences.

### 2.2. Simulation Studies

In order to examine the performance of the GLM in modeling the dopamine displacement, we generated four simulations with varying task paradigms and noise levels by solving the differential equation of the LSSRM (Eq. 2), and compared the performance of the GLM with the performance of the operational equation of the LSSRM. We simulated the actual task data with a temporal resolution of 30 s and scan length of 50 mins, in which each task block lasted 10 minutes. The ode23s toolbox in Matlab was used for solving the differential equation in LSSRM. C(t) was simulated with R = 0.95, k_2_ = 0.23, BP = 1.86, as estimated from real data. The reference curve *C_r_*(*t*) was extracted from the cerebellum and was fitted with a rational function to eliminate noise:

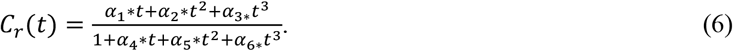

To mimic the real situation, noise proportional to the radioactivity concentration was added to the noise-free C(t) as:

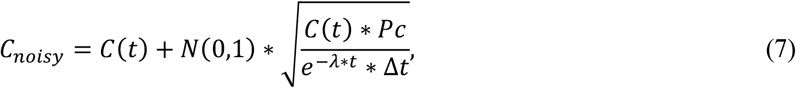

where the exponential term was used to remove the decay correction. We used a scaling factor *Pc* to control the noise level, *N*(0,1) denotes a Gaussian distribution with zero mean and unit variance, and *λ* is 0.034 for ^11^C.

Although the statistics of nuclear decay follow the binomial law, and event detection with PET detectors can be well fitted with a Poisson distribution, a Gaussian distribution is more effective for simulating reconstructed PET images considering several additive sources of error (Logan et al., 2001).

To examine different aspects of the proposed model, we performed four simulations as follows.

**<u>Simulation 1:</u>** One single task block and fixed noise levels. The activation level *r* in Eq. 2 was simulated 1000 times to vary between 0 and 0.1 to match observations from real data. The noise level was fixed with *Pc* = 4. A TAC example is shown in Fig. 1A along with the modelling of LSSRM and GLM.

**Fig. 1.**
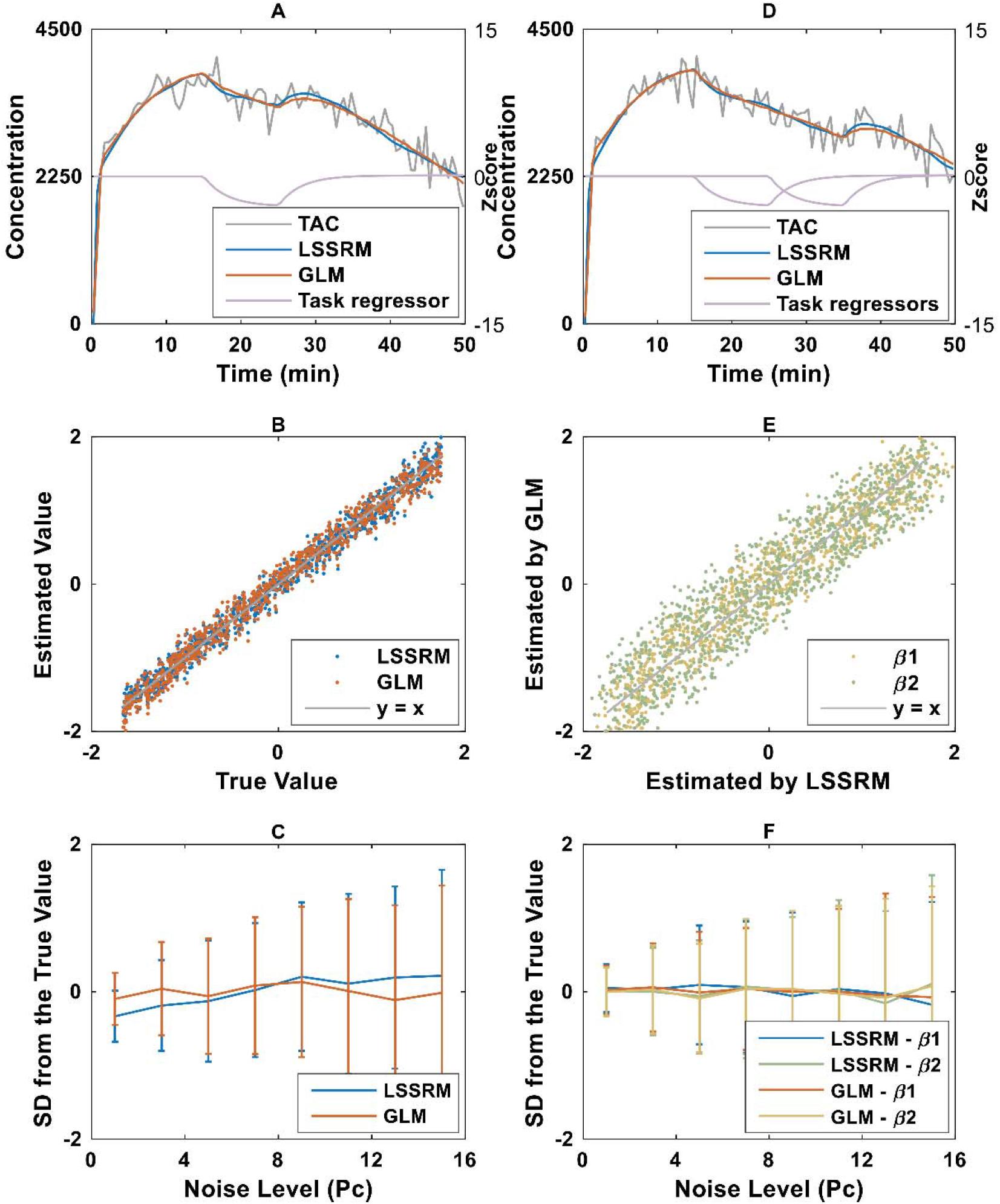
Examples and results of simulation studies, four simulations with varying task paradigms and noise levels were generated for the performance comparison of the general linear model (GLM) and the linear extension of the simplified reference region model (LSSRM). (A) An example of time activity curve (TAC, grey) simulated with a single monetary incentive delay (MID) task, the TACs fitted by the LSSRM (blue) and the GLM (orange), and the normalized task regressor (Z scores) used in GLM (purple); (B) The estimated and true values in the Simulation 1 (one single task block and fixed noise levels), there is a high consistency between the values estimated by LSSRM/GLM and the true values; (C) The estimated values of LSSRM and GLM under different noise levels in Simulation 2 (one single task block and random noise levels), greater bias (standard deviation, SD) was observed under higher noise level for both approaches; (D) A TAC example simulated with two MID tasks in a single scan (grey), the TACs fitted by the LSSRM (blue), the GLM (orange) and the normalized task regressors in GLM (purple); (E) The estimated values in the Simulation 3 (two task blocks and fixed noise levels) through the LSSRM and the GLM, the estimated values from the two models showed high correlations for both tasks (β1 and β2); (F) The estimated values of LSSRM and GLM under different noise levels in Simulation 4 (two task blocks and random noise levels), similar to the Simulation 2, an increased SD was shown with heightened noise level for both tasks (β1 and β2 respectively).

**<u>Simulation 2:</u>** One single task block and random noise levels. The *r* was fixed to 0.04 with noise level *Pc* varying between 0 and 16 for 1000 times to test the robustness of the two models to noise disturbance.

**<u>Simulation 3:</u>** Two task blocks and fixed noise levels. To mimic the experimental design in the human study, two MID tasks were implemented with the second one beginning directly after the end of the first task, *r* for each task was randomized between 0 and 0.1 for 1000 times with *Pc* = 4. A TAC example is shown in Fig. 1D along with the modelling of LSSRM and GLM.

**<u>Simulation 4:</u>** Two task blocks and random noise levels. Activation levels for the tasks were set to *r*1 = 0.04, *r*2 = 0.03, and *Pc* varied between 0 and 16 for 1000 times.

All simulation data were modelled through the classic LSSRM, and the parameters were estimated using the proposed GLM, with both ordinary least squares (LS) analysis and the weighted linear least squares (WLS) analysis (the weighting parameter 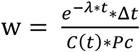). The Akaike information criterion (AIC) was applied to evaluate the performance of the different statistical models, with a lower AIC indicating higher goodness of fit. The Pearson correlation coefficient *(r_p_*) and the intraclass correlation coefficient (ICC) of the estimated values and the true values were calculated to evaluate the reliability of each method. In addition, the simulation results of Simulation 1 were mapped onto a striatum template (Tzourio-Mazoyer et al., 2002) for a voxel-by-voxel visual comparison of the findings of GLM and LSSRM.

## 3. Human Study

### 3.1. Participants

Fifteen MDD participants (13 unmedicated; 8 female; mean age = 32. 2 years) and fourteen CTL subjects (all unmedicated; 10 female; mean age = 32.5 years) were recruited for the study. No participants had a history of head injury or drug abuse within six months prior to the experiment. Depressed participants met diagnosis criteria for MDD based on Structured Clinical Interview for DSM-5. All participants completed the 10-item ruminative response scale (RRS) (Treynor et al., 2003), which includes two subscales: brooding and reflection. A brief summary of participants’ demographic information and rumination trait scores are shown in Table 1. More details about the participants can be found in a previous publication (Hamilton et al., 2018). The study protocol was approved by the Stanford University Institutional Review Board and maintained compliance with federal, state, and local regulations on medical research (IND 123, 410). Written informed consent was obtained from all participants.

**Table 1.**
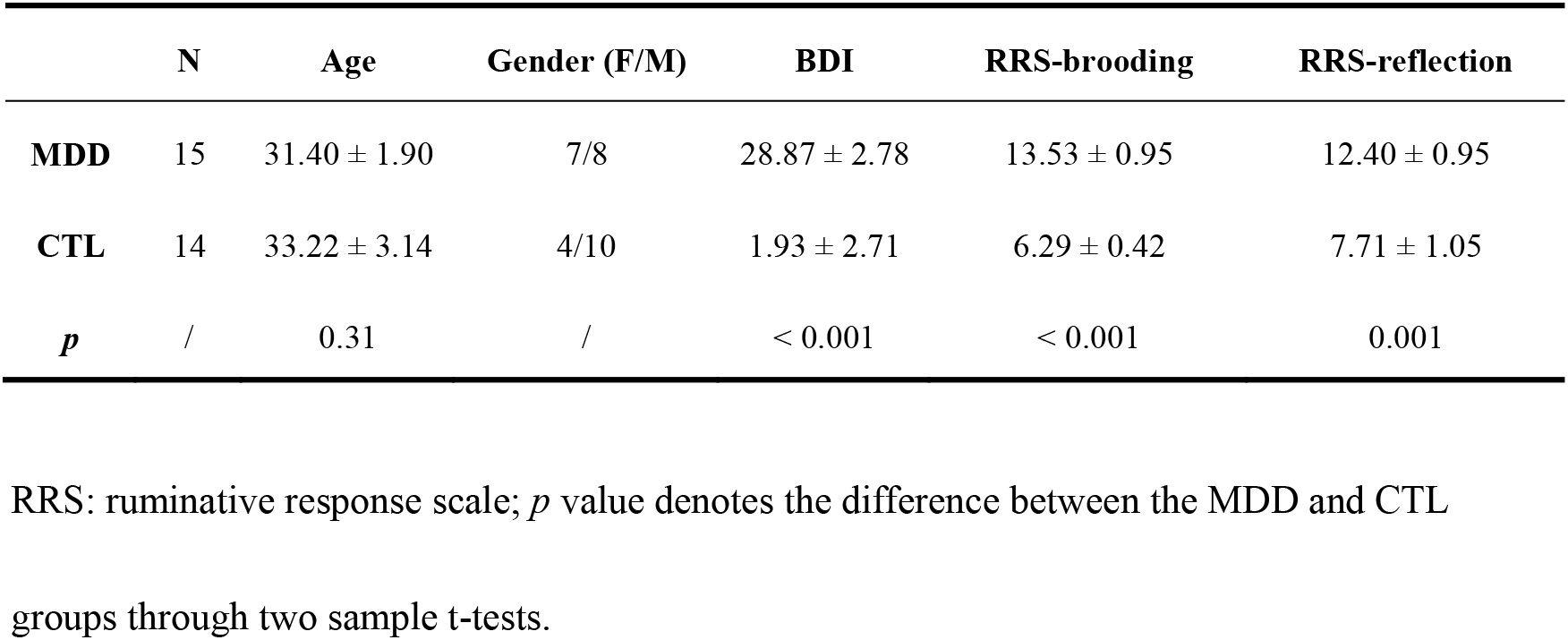
The demographics and rumination scores of subjects.

|  | **N** | **Age** | **Gender (F/M)** | **BDI** | **RRS-brooding** | **RRS-reflection** |
| --- | --- | --- | --- | --- | --- | --- |
| **MDD** | 15 | 31.40 ± 1.90 | 7/8 | 28.87 ± 2.78 | 13.53 ± 0.95 | 12.40 ± 0.95 |
| **CTL** | 14 | 33.22 ± 3.14 | 4/10 | 1.93 ± 2.71 | 6.29 ± 0.42 | 7.71 ± 1.05 |
| ***p*** | / | 0.31 | / | < 0.001 | < 0.001 | 0.001 |

RRS: ruminative response scale; *p* value denotes the difference between the MDD and CTL
groups through two sample t-tests.

### 3.2. Data Acquisition

All participants underwent a simultaneous fPET-fMRI examination on a time-of-flight (TOF) PET-MRI scanner (3T SIGNA PET-MRI; GE Medical Systems, USA). The fMRI images were acquired axially using a spiral-in/out (Glover and Law, 2001) imaging sequence with TR = 2500 ms, TE = 30 ms, FOV = 22.0 cm^2^, Matrix = 64 × 64, 27 slices, and voxel size = 3.44 × 3.44 × 3.5 mm^3^. PET images were dynamically reconstructed by an ordered subsets expectation maximization (Hudson and Larkin, 1994) algorithm (28 subsets, 3 iterations) and resampled to match the spatial resolution of fMRI images with TR = 30 s. Scatter correction, MR-based attenuation correction, point spread function correction, dead time correction, and decay correction were performed during reconstruction using the scanner’s built-in software.

### 3.3. Experimental Design

The scanning procedure is shown in Fig. 2A. PET scanning took place for the entire 42-minute /50-minute scanning session. The radiosynthesis and injection procedures are described in the previous publication (Hamilton et al., 2018). In general, the synthesized [^11^C] Raclopride was injected via the left antecubital vein with 10 mCi over 60 s, and the overall radioactivity was 10.6 ± 3.9 Ci/μmol for all subjects. The bolus injection of [^11^C] Raclopride was administered one minute after the PET scan started (the fPET time series reconstruction was initiated when 4000 counts per second were reached), and we started a 32-minute-long fMRI scan 9 mins after the injection to allow for the uptake of the radiotracer. The fMRI scan included two resting-state acquisitions before and after the two reward tasks.

**Fig. 2.**
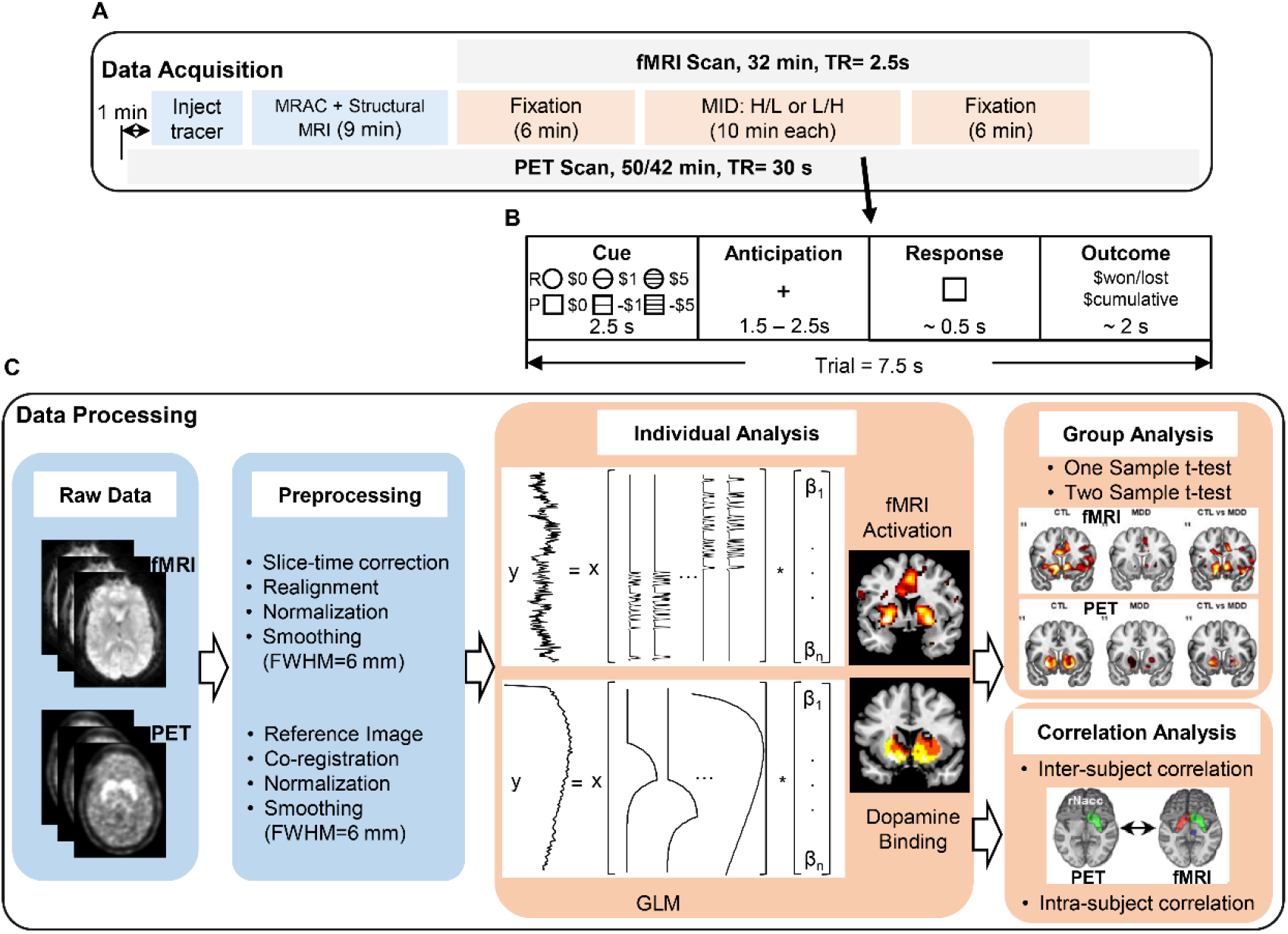
The scanning procedure, the experimental design and the data analysis. (A) The scanning procedure for PET and fMRI imaging. MRAC: MRI-based attenuation correction, MRI: magnetic resonance imaging, H/L: high-stake to low-stake order, L/H: low-stake to high-stake order; (B) A trial illustration for the monetary incentive delay (MID) task, each task consists of the cue, anticipation, response and the outcome periods. R: reward; P: punish; (C) Data analyses for the task fMRI and fPET data.

The MID task, a well-developed reward paradigm, was employed to induce reward processing. During the task scan, participants performed two MID tasks (10-min each) at different stake levels (low: ±$1; high: ±$5) in a counterbalanced sequence of either low-stake to high-stake order (LH) or high-stake to low-stake order (HL). The trial structure for the task is depicted in Fig. 2B, with each trial lasting 7.5s. At the beginning of each trial, a cue was presented indicating the possibility of gain (circle) or loss (square) in this trial; horizontal lines within the circle/square denoted the amount of money for the current trial (no line for no-gain/loss trials, 1 line for $1 and 3 lines for $5). After ~ 2 s anticipation with a “+” shown on the screen, a square was presented in the response period and subjects were asked to press the button as soon as possible, to win money or to avoid losing money. Finally, the current winnings/losses and the total money accumulated were shown on the screen for ~ 2 s. Each task consisted of 80 trials, with 21 gain trials and 20 no-gain trials, 18 loss trials and 21 no-loss trials in the low-stake task, and 19 gain trials and 21 no-gain trials, 20 loss trials and 20 no-loss trials in the high-stake task. Subjects were given $10 at the beginning of the scan. To increase the uncertainty of money accumulation, the task difficulty level was automatically adjusted by modulating the target stimulus duration to keep the subjects’ accuracy to around 67%. We were primarily interested in the anticipation period since it has been reported to evoke the strongest activation in the reward system during reward processing (Knutson et al., 2001; O’Doherty et al., 2002).

### 3.4. Data Analysis

Data processing was implemented in DPABI4.4 (http://rfmri.org/DPARSF), SPM8 (http://www.fil.ion.ucl.ac.uk/spm/software/spm8/) and self-developed MATLAB code. Our aims were to compare the fMRI activation and fPET activation between the MDD and CTL, and to explore the coupling between the dopaminergic system and the reward network during reward processing (MID task). To this end, three types of metrics were explored and compared between the MDD and CTL participants: (1) fMRI activation during MID tasks; (2) fPET activation during MID tasks; and (3) the inter- and intra-subject correlation between the two modalities (fMRI activation and dopamine release). The analysis pipeline is outlined in Fig. 2C.

#### 3.4.1. fMRI Data Analysis

A routine preprocessing pipeline was applied to the fMRI data, including slice-time correction, realignment, normalization to standard (MNI) space, and spatial smoothing (FWHM=6 mm). Percentage signal change was calculated with respect to the mean activity over the entire task session. Preprocessed data were entered into the GLM at the individual level with each task condition modelled as one covariate: anticipation, response, outcome for low/high reward/no-reward/punishment/no-punishment conditions. We should note that our task has a relatively long cue period (2.5 s); consequently, participants could start to anticipate right after seeing the cue. Therefore, we included the cue period in the anticipation condition to increase activation detection. Twenty-four motion parameters (Friston et al., 1996) were included in the GLM as covariates of no interest. The activation maps of anticipation of low/high stake reward/punishment versus no-reward/no-punishment contrasts were estimated for correlation analysis with the concurrent PET measurements of dopamine release.

#### 3.4.2. fPET Data Analysis

The fPET data concurrently collected with fMRI acquisitions (10-41 minutes) were first realigned based on the motion parameters estimated from fMRI data. Next, we averaged the 2-8 minutes pre-task fPET data to generate a reference image that contained both structural outline and binding information for a rigid body realignment across different time points and then co-registered all fPET images to the mean fMRI image. fPET images were normalized and smoothed with the same parameters as for fMRI. After preprocessing, fPET data were modelled by the GLM as described in the simulation study (Eq. 5) to detect the level of task-induced dopamine release. The fPET task data has been analyzed through LSSRM in our previous study (Hamilton et al., 2018), however, no significant binding potential difference of the MDD and the CTL group was observed, we would expect that modelling through GLM might distinguish the two groups. Given the coarser temporal resolution of fPET relative to fMRI, and corresponding reduction in the number of fPET acquisitions, we did not include the movement parameters in the GLM to preserve the degrees of freedom in the model. The intrinsic resolution of PET images is 6.4 mm^3^ (estimated by the autocorrelation of noise maps), alleviating the potential concern of slight head motion.

#### 3.4.3. Group Analysis

For both fMRI and fPET analyses, a one sample t-test was conducted to evaluate the group-level fMRI activation and dopamine release, and a two sample t-test was conducted to compare the results between the MDD and CTL groups. The voxel-wise neural activation and the dopamine release (β values fitted in the GLM) were correlated with the rumination level, respectively. The statistical significance criterion was set at *p* < 0.05, FDR-corrected for multiple comparisons.

##### Inter-subject Correlation Analysis

Dopamine binding in the bilateral NAcc (β values in PET signal-based GLM analysis) was extracted from the fMRI activation maps (Fig. 5), and correlated with the levels of whole-brain fMRI activations (β values in fMRI signal-based GLM activation) for the CTL and MDD groups separately. The inter-subject correlation coefficients of the two groups were compared through a permutation test using 500 randomizations (Nichols and Holmes, 2002).

##### Intra-subject Cross-modal Correlation Analysis

To assess the coupling of the fPET activation (dopamine binding) and the fMRI activation at the individual level, we correlated the voxel-wise fPET activation with the corresponding fMRI activation for each subject in the striatum, and the group difference between the CTL and the MDD was assessed. To increase the SNR of the input data for the correlation analysis, we measured region of interest (ROI)-based correlation instead of voxel-based correlation by dividing the striatum into 8 ROIs, including bilateral NAcc, caudate, putamen and pallidum following the Harvard Oxford Atlas (Fig. 6A). FMRI activation and the dopamine release of those ROIs were correlated to evaluate the coupling relation within each individual. The intra-subject cross-modal correlation was correlated with the rumination levels with *p* < 0.05 as the threshold.

## 4. Results

### 4.1. Simulation Studies

**Simulation 1** (one single task block and fixed noise levels): The dopamine release estimated through the LSSRM and GLM is shown in Fig. 1B: there was a very high consistency between the values measured through both methods and the true values (LSSRM: *r_p_* = 0.9918, ICC = 0.9959; GLM: *r_p_* = 0.9899, ICC = 0.9949). The AIC values of the 1000 simulations were significantly lower in the GLM than the LSSRM (paired t-test, *p* < 0.001, Fig. S2A), suggesting a better fitting using the GLM. Besides, the AIC values were reduced using the WLS analysis; however for the detection of dopamine release (T value), the correlation and ICC values with true values from WLS did not improve (Table S1), so the LS was employed in the subsequent analyses. The simulated dopamine release using the LSSRM and GLM in a striatum template is shown in Fig. S3.

**Simulation 2** (one single task block and random noise levels): As expected, with higher noise level simulated, there was a more significant bias of the estimated values from the true values for both methods (Fig. 1C). Although the *r_p_* and ICC with the true value were low for both methods (< 0.1), there was a relatively high consistency of the estimated values from the LSSRM and the GLM (*r_p_* = 0.7605, ICC = 0.8640), the AIC values of the 1000 simulations were significantly lower in the GLM than the LSSRM (paired t-test, *p* < 0.001, Fig. S2B).

**Simulation 3** (two task blocks and fixed noise levels): Similar to Simulation 1, we observed a high reliability of the simulated values from both methods for two tasks (Fig. 1E, LSSRM: *r*_*p_task*1_ = 0.9905, *r*_*p_task*2_ = 0.9880, *ICC*_*task*1_ = 0.9952, *ICC*_*task*2_ = 0.9940; GLM: *r*_*p_task*1_ = 0.9504, *r*_*p_task*2_ = 0.9304, *ICC*_*task*1_ = 0.9746, *ICC*_*task*2_ = 0.9640), the correlation and ICC values of the LSSRM were slightly higher than the GLM, however the AIC values of the 1000 simulations were significantly higher in the LSSRM than the GLM (paired t-test, *p* < 0.001, Fig. S2C).

**Simulation 4** (two tasks and random noise levels): Similar to Simulation 2, the estimated values lost accuracy with higher noise levels for both tasks (Fig. 1F). There was a relatively high consistency of the estimated values from the LSSRM and the GLM (*r*_*p_task*1_ = 0.7266, *r*_*p_task*2_ = 0.7560; *ICC*_*task*1_ = 0.8417, *ICC*_*task*2_ = 0.8610), similar to previous simulations, the AIC values of the 1000 simulations were significantly lower in the GLM than the LSSRM (paired t-test, *p* < 0.001, Fig. S2D).

Thus, in all simulations, the AIC values were significantly lower for the GLM method, suggesting a more precise modeling of the TACs using the GLM.

### 4.2. Human Study

#### 4.2.1. FMRI Activation during the Reward Anticipation and Its Correlation with Rumination

The high-reward versus no-reward anticipation contrast induced the most significant activation among all conditions and is reported in Fig. 3; the findings of high-punishment versus no-punishment are shown in the supplementary material (Fig. S4). The anticipation of high-reward versus no-reward induced activation in the striatum, including the putamen, the caudate, and the NAcc (Fig. 3A and Fig. 3B) in both groups. In addition, the VTA, bilateral thalamus, bilateral visual cortex, bilateral anterior insula and sensory motor areas were also activated (Fig. 3A and Fig. 3B). Group analysis revealed an overall reduced activation in MDD (Fig. 3B) compared with CTL (Fig. 3A) in the NAcc, the ventral medial prefrontal cortex (vmPFC) and the posterior cingulate cortex (PCC), etc. (Fig. 3C). The activation of anticipation under high-reward versus noreward condition in the right middle putamen was significantly correlated with the reflection subscale of RRS in the CTL (*r_p_* = −0.61, *p* = 0.01, Fig. 3D) and the MDD group (*r_p_* = −0.79, *p* < 0.001, Fig. 3E). In addition, the activation in the vmPFC (*r_p_* = −0.79, *p* < 0.001, Fig. 3F) was also correlated with the reflection subscale in the MDD group.

**Fig. 3.**
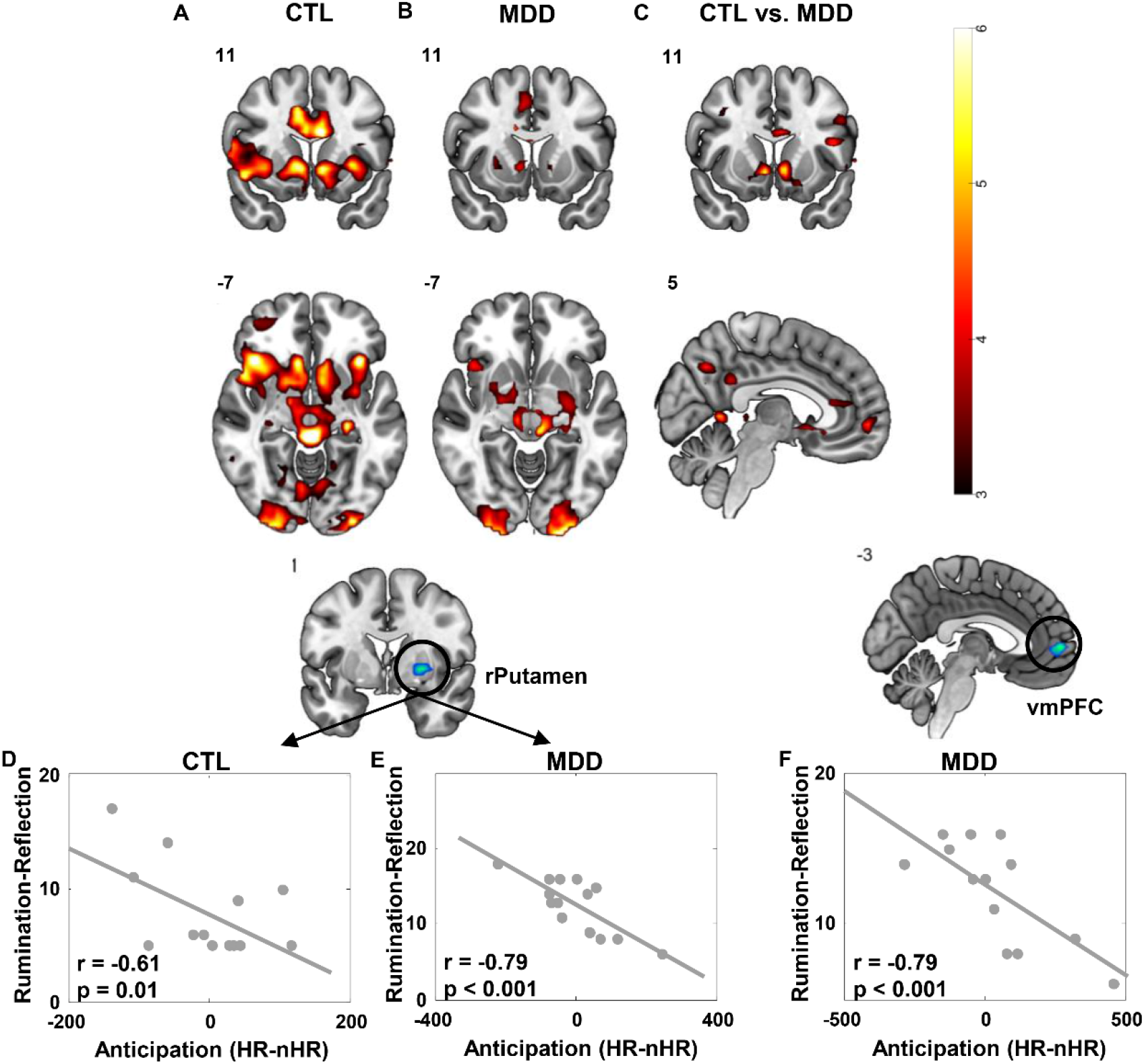
The fMRI activations under high reward monetary incentive delay (MID) tasks and the correlation with reflection rumination scores. High reward ($5) versus no reward anticipation contrasts in CTL (A), MDD (B), and CTL versus MDD (C). The fMRI activation of the right middle putamen in the CTL (D), the fMRI activation of the right middle putamen (E) and the fMRI activation of the vmPFC (F) in the MDD were negatively correlated with the reflection rumination scores.

#### 4.2.2. Dopamine Release during the Reward Task and Its Association with Rumination

During the MID task, increased dopamine binding in the striatum, including the bilateral NAcc, was observed in both the CTL (Fig. 4A) and MDD (Fig. 4B) groups, and the activation was greater in the CTL group in the right anterior caudate and right middle putamen (Fig. 4C), which is consistent with the fMRI findings. We also observed a correlation of the dopamine release in the left putamen (*r_p_* = −0.67, *p* = 0.006, Fig. 4D) and the right NAcc (rNAcc, *r_p_* = −0.69, *p* = 0.004, Fig. 4E) and the reflection subscale in the MDD group.

**Fig. 4.**
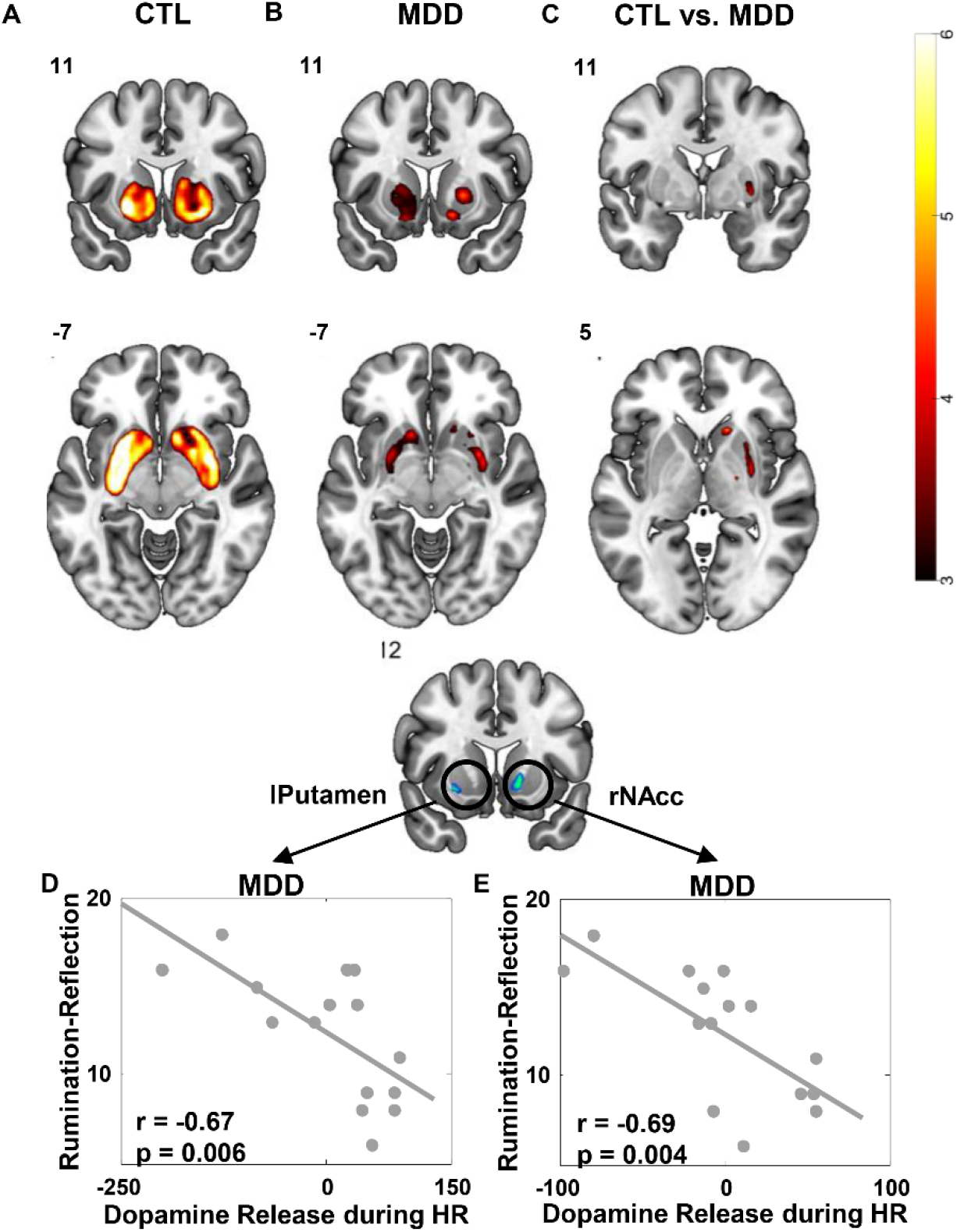
The Dopamine release during the monetary incentive delay (MID) task and the correlation with reflection rumination scores. Dopamine activation during the MID task in CTL (A), MDD (B), and CTL versus MDD (C). The dopamine release in the left putamen (D) and the right nucleus accumbens (rNAcc, E) was negatively correlated with the reflection rumination scores in the MDD.

#### 4.2.3. Inter-subject Correlation between the Dopamine Release in the rNAcc and the fMRI Activation

The inter-subject correlation coefficients was significantly higher in the CTL group in the bilateral NAcc, VTA, bilateral anterior insula, and the anterior cingulate cortex, etc. (Fig. 5A). For the CTL group, the relationship between dopamine binding in the rNAcc and fMRI activation in the VTA (*r_p_* = 0.80, *p* < 0.001, Fig. 5B) and in the bilateral NAcc (left NAcc: *r_p_* = 0.70, *p* = 0.003, Fig. 5C; right NAcc: *r_p_* = 0.66, *p* = 0.007, Fig. 5D) are shown in Fig. 5. However, no significant correlation was observed for the MDD group.

**Fig. 5.**
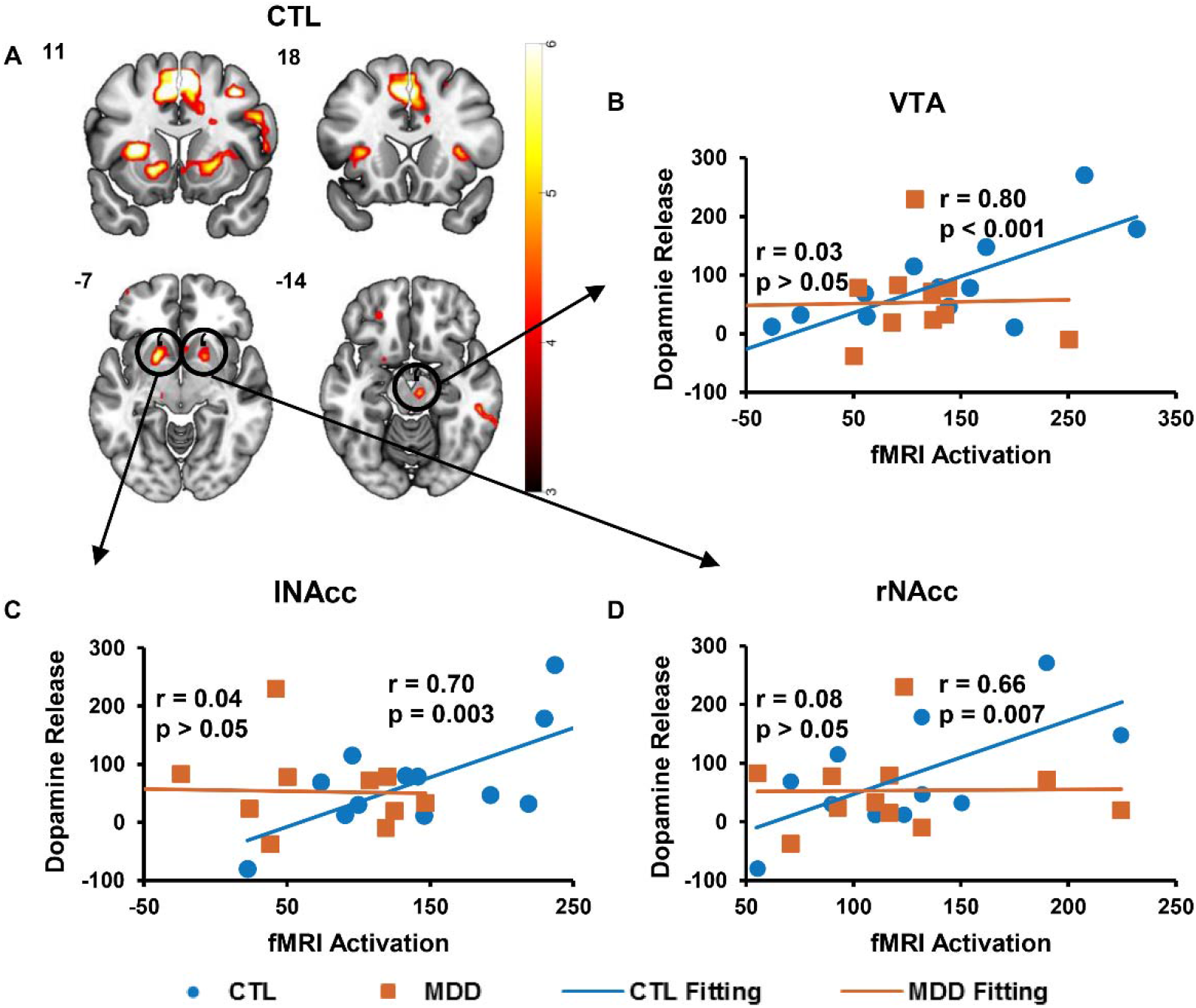
Inter-subject correlation of the dopamine release in the right nucleus accumbens (rNAcc) and the whole-brain fMRI activations. (A) Voxel-based analysis showed significant higher correlation of dopamine displacement in rNAcc and neural activity of the ventral tegmental area (VTA, B), bilateral NAcc (C, D) and bilateral anterior insula in the CTL group, while the MDD group showed no correlation.

#### 4.2.4. ROI-to-ROI Intra-subject Correlation between the Dopamine Release and the fMRI Activation

In the CTL group, most participants had a significant correlation between their neural activation and dopamine level in an ROI-to-ROI-based cross-modal comparison (Fig. 6B); one outlier (< mean – 3 standard deviations) was excluded from the analysis. One-sample t-test yielded a significant intra-subject correlation at the group level in the CTL group (*p* = 0.003) but not in the MDD group; a two-sample t-test yielded a significantly lower correlation in the MDD group than the CTL group (Fig. 6C, *p* = 0.02). Notably, the intra-subject correlation of the fMRI activation and the dopamine release in the CTL was negatively associated with reflection rumination scores (*r_p_* = −0.69, *p* = 0.02).

**Fig. 6.**
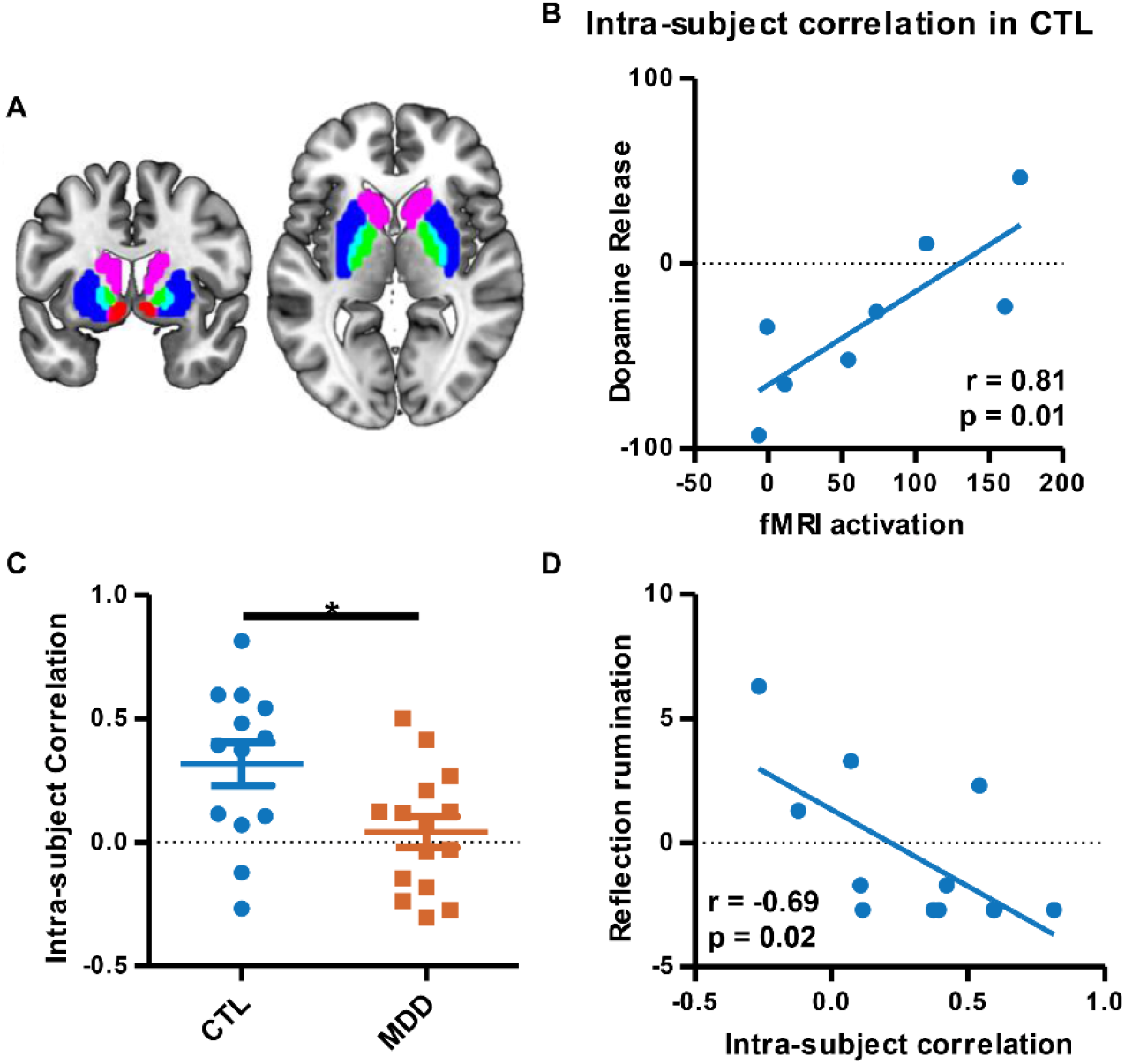
Intra-subject correlation of the fMRI activations and the dopamine release in the striatum under high-stake reward tasks. (A) ROIs used in the correlation analysis, including the nucleus accumbens (NAcc, red), caudate (violet), putamen (blue) and pallidum (green); (B) An example of the significant correlation between fMRI activations and the dopamine release in the CTL; (C) The intra-subject correlation in the CTL and MDD; (D) The fMRI-PET intra-subject correlation in the CTL was negatively correlated with the reflection rumination score. *: p < 0.05

## 5. Discussion and Conclusion

The present study used a concurrent fPET-fMRI approach to examine reward processing during a MID task in CTL and MDD participants and examined the relation between dopamine release/binding inferred from [^11^C] Raclopride fPET and neural activity indexed by BOLD responses in the reward system, the impact of trait rumination on the reward processing was assessed by correlating it with the dopamine release, fMRI activation and their coupling. The feasibility of using GLM in estimation dopamine release during reward task was also explored through simulation tasks. Overall, decreased fMRI activation and dopamine release/binding were observed in the MDD group, indicating the reduced capability of reward processing in MDD. In the MDD group, individuals with lower fMRI activations in the right middle putamen and the vmPFC had higher reflection rumination scores, and individuals with lower dopamine release/binding in the left putamen and the right NAcc also showed higher reflection rumination scores, these findings suggest that the deficiency in the reward network and the dopaminergic system might couple with individuals’ rumination levels. As we have hypothesized, significant inter-subject and intra-subject correlation between the dopamine release/binding and the fMRI activation was only observed in the CTL group, indicating that the disconnection between dopaminergic system and striatum may be a critical factor in the development of MDD.

FMRI activation of the NAcc was lower in the MDD group than in the CTL group, consistent with previous studies (Pizzagalli et al., 2009; Takamura et al., 2017; Ubl et al., 2015a). The striatum, especially the NAcc, is a core node of the reward system, and its activity is related to reward information from external stimuli and individual goal-oriented behavior (Kohls et al., 2012). Specifically, the striatum is thought to bridge the limbic and motor systems, translating rewarding stimulation in the external environment into the motivation of “wanting” that drives behavior (Kohls et al., 2012). “Wanting” is a process of categorizing external stimuli into attractive and urgent events. The decreased striatum activity observed here might indicate the weakening of this driving force, and its relevance to the reflection rumination further supports that reduced activation of the reward system may cause the individual to reduce focusing on the external stimuli and pay more attention to the self-related events. The vmPFC is a core node of default mode network and is associated with numerous complex functions such as autonomic and endocrine regulation, emotion, emotion regulation, semantic memory and self-directed cognition (Roy et al., 2012). Resting-state fMRI studies have reported increased functional connectivity of the vmPFC and the orbital frontal cortex and its correlation with levels of rumination (Hamilton et al., 2011). Moreover, vmPFC activity is associated with individuals’ preference for particular stimuli; that is, high-value stimuli are more likely to activate vmPFC than low-value stimuli (Plassmann et al., 2008). Lower vmPFC activation in MDD reflects that MDD has a higher threshold for stimuli that can be of interest, which may be the neural basis of anhedonia, a primary symptom of MDD characterized by a reduced capacity to experience pleasure.

We found that striatal dopamine release/binding was lower in the MDD group than the CTL group during the MID task, and that dopamine release in the left putamen and the right NAcc was negatively associated with reflection rumination, which is consistent with the findings related to fMRI. Previous studies on baseline dopamine receptor binding have reported inconsistent results (D’Haenen H and Bossuyt, 1994; de Kwaasteniet et al., 2014; Ebert et al., 1996; Hamilton et al., 2018; Hirvonen et al., 2008b; Klimke et al., 1999; Meyer et al., 2006; Montgomery et al., 2007b; Parsey et al., 2001; Pecina et al., 2017; Shah et al., 1997; Yang et al., 2008). Of note, our previous publication using the current dataset showed increased *baseline* binding potential in the MDD (Hamilton et al., 2018), but showed no difference in the *task* binding potential. In the current study, we used the GLM instead of the kinetic modeling in analyzing the dynamic fPET data and detected a decrease of dopamine release in the MDD group, our findings of decreased dopamine release in MDD suggest that the dopaminergic system activity triggered by the reward process can be utilized to distinguish between MDD and CTL.

Dopaminergic activation of the right NAcc was found to correlate with fMRI-measured neural activation in bilateral NAcc, VTA and bilateral anterior insula, underscoring the importance of the interaction between the dopaminergic system and the reward network for the normal functioning during the reward processing. We failed to observe an interaction between the two systems in the MDD group, suggesting that the reduction in activation in the reward system may be due to the reduced dopamine level or the failure of dopamine to bind well to the target receptors to drive the reward system. Moreover, we observed an fPET-fMRI intra-subject correlation of the fMRI activation and the dopamine release in the CTL and the correlation was negatively associated with the reflection rumination scores, which suggests that the coupling of the reward system and the dopaminergic system could be another imaging marker for the rumination level.

We should note that this study found a link between the reward response and the reflection rumination score, rather than the brooding rumination scale. Although the brooding subscale has been frequently reported to be more relevant to the symptoms of depression (Takano and Tanno, 2009), the reflection rumination has also been reported to associate with depressive symptoms, and reflection scores have been previously reported to positively correlate with suicidal thoughts one year later (Miranda and Nolen-Hoeksema, 2007).

In this study, we also demonstrated the feasibility of using the GLM to analyze dynamic Raclopride fPET signals. The simulation results showed improved performance of GLM with the traditional kinetic modeling in detecting the dynamic dopamine release, without invoking steadystate assumptions and the selection of a suitable reference region. Since the GLM is simple and widely available in fMRI analysis toolboxes, we suggest it may be an effective alternative to the traditional methods.

fMRI data acquisition was initiated before the MID tasks commenced (Fig. 2A) for two reasons: (1) to allow tracer wash into the brain region to complete; and (2) to enable any transient interactions between starting the fMRI gradients and PET detectors to subside (eddy currents induced in the PET detectors by the gradients could cause transiently-uncompensated temperature changes, leading to sensitivity changes). Such effects were small in our system, however (Deller et al., 2018).

The current research has several limitations. First, the cue period in the MID task was longer than typical (2.5 seconds), which could be shortened in future studies. Second, the gold standard data used in the simulation experiment was generated by kinetic modeling, so there may be a bias in comparing the performance of the GLM and the kinetic modeling in this context. In the future, it would be useful to consider less confounded simulation experiments to compare the two methods. Finally, the sample size of both the CTL group and the MDD group is small, and the results of this study need to be replicated in larger samples.

Overall, we examined the coupling of the dopaminergic system and the reward network in MDD and CTL by conducting a simultaneous [^11^C] Raclopride fPET-fMRI study. Our findings suggest that concurrent fPET and fMRI may provide valuable information in understanding the pathophysiology of psychiatric disorders, e.g., MDD examined here. We note that concurrent acquisition of the two modalities alleviates concerns that differences in the mental state between separately acquired fPET and fMRI data may diminish the reliability of their combination. Finally we suggest that dynamic fPET data may be analyzed using a GLM with an exponential tracer response function that is analogous to fMRI analyses using a hemodynamic response function to model BOLD contrast.

## Acknowledgements

We thank Brian Knutson for assistance with the task design, Emily Livermore, Christina Schreiner, Maddie Pollak, Monica Ellwood-Lowe, Sophie Schouboe, and Moema Gondim for their help with participant recruitment and data collection, Dawn Holley and Hershel Mehta for their support for scanning, Fred Chin and Bin Shen for radiosynthesis preparation, Lihong Wang for valuable discussions. This work was supported in part by NIH EB015891 (XZ, FW, GHG, JC) and Weston Havens Foundation (IHG, JPH, MDS).

