## Supplementary Materials for "Decoupling of Dopamine Release and Neural Activity in Major Depressive Disorder during Reward Processing Assessed by Simultaneous fPET-fMRI"

**Supplementary Material**

Table S1 Performance comparison of the least square (LS) and the weighted least square (WLS) analysis in Simulation 1

|  | **AIC** | **Dopamine release**  **(T value)** | **Correlation with true value** | **ICC with true value** |
| --- | --- | --- | --- | --- |
| **GLM-LS** | 1351.50 ± 0.49 | 6.22 ± 0.11 | 0.9899 | 0.9949 |
| **GLM-WLS** | 110.14 ± 0.0002 | 5.93 ± 0.11 | 0.9896 | 0.9948 |
| **p** | < 0.001 | < 0.001 | / | / |
| **LSSRM-LS** | 1368.70 ± 0.40 | 6.59 ± 0.12 | 0.9918 | 0.9959 |
| **LSSRM-WLS** | 110.15 ± 0.0001 | 6.36 ± 0.12 | 0.9914 | 0.9957 |
| **p** | < 0.001 | < 0.001 | / | / |

AIC: Akaike information criterion; p value denotes the difference between the LS and WLS through paired t-test.

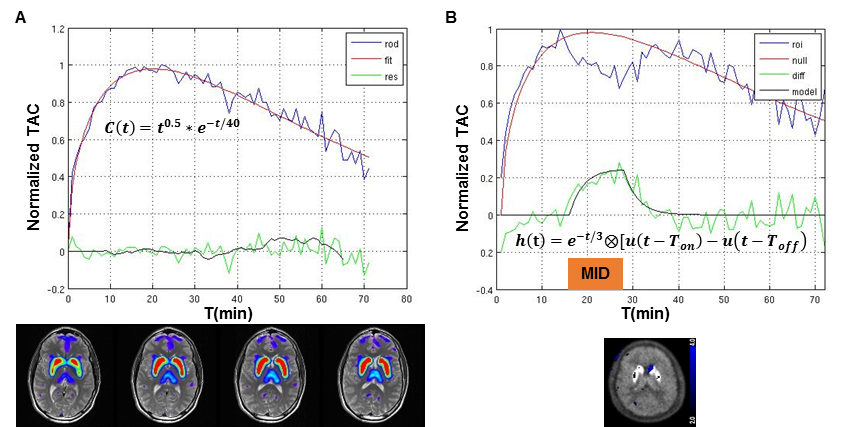

Fig. S1 The normalized baseline and task time activity curves (TACs) in the pilot study. (A) The baseline TAC (‘rod’) of the striatum (red, bottom panel) where the dopamine D2 receptor locates, the fitted TAC by GLM (‘fit’) and the residuals (‘res’); (B) The TAC (‘roi’) of the dorsal putamen (blue, bottom panel) during a monetary incentive delay (MID) task. After regressing out the baseline uptake and wash-out curve (‘null’) derived from the baseline scan, the residuals (‘diff’) were fitted well by an impulse response function of the exponential decay function convolved by the task design (‘model’).

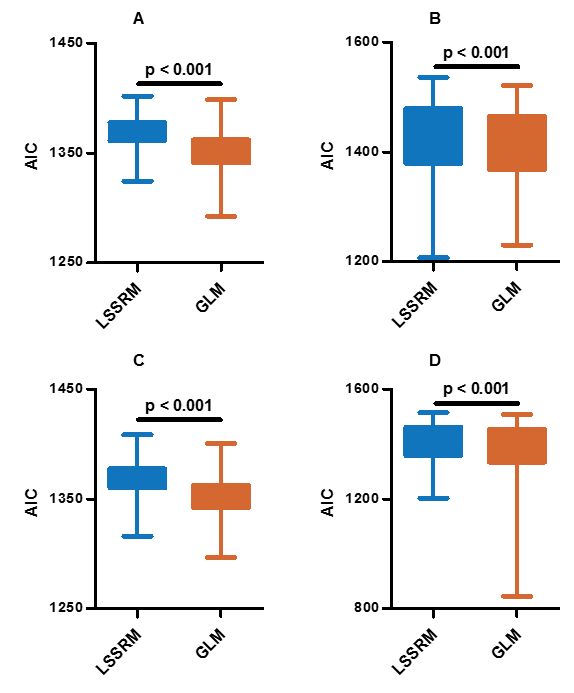

Fig. S2 The Akaike information criterion (AIC) values of the 4 simulations. (A) Simulation 1: one single task block and fixed noise levels; (B) Simulation 2: one single task block and random noise levels; (C) Simulation 3: two task blocks and fixed noise levels; (D) Simulation 4: two task blocks and random noise levels.

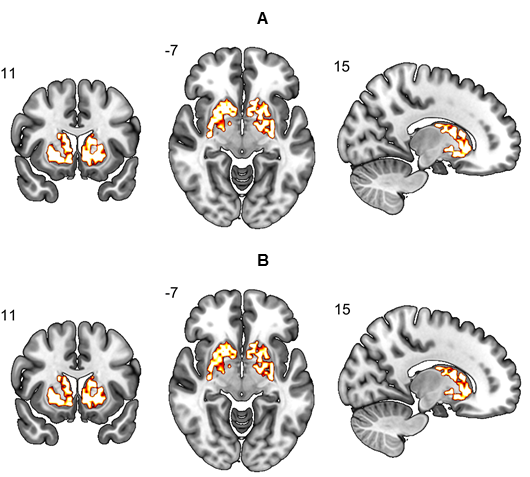

Fig. S3 The simulated dopamine release in the striatum template. (A) Dopamine release estimated by the linear extension of the simplified reference region model (LSSRM, orange); (B) Dopamine release estimated by the general linear model (GLM, orange).

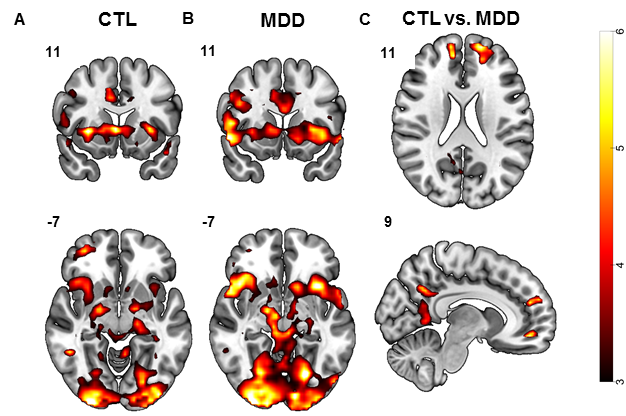

Fig. S4 The fMRI activation of high punishment ($5) versus no punishment anticipation contrasts in CTL (A), MDD (B), and CTL versus MDD (C).
